# Mechanotransduction and inflammation: an updated comprehensive representation

**DOI:** 10.1101/2024.06.12.598454

**Authors:** Vennila Suryiagandhi, Ying Ma, Veronica Paparozzi, Tiziana Guarnieri, Biagio di Pietro, Giovanna Maria Dimitri, Paolo Tieri, Claudia Sala, Darong Lai, Christine Nardini

## Abstract

Mechanotransduction is the process that enables the conversion of mechanical cues into biochemical signaling. While all our cells are well known to be sensitive to such stimuli, the details of the systemic interaction between mechanical input and inflammation are not always well integrated. Often, they are considered and studied in relatively compartmentalized areas, and we therefore argue here that to understand the relationship of mechanical stimuli with inflammation – with high translational potential - it is crucial to offer and analyze a unified view of mechanotransduction. We therefore offer here a pathway representation, recollected with the standard systems biology markup language (SBML) and explored with network biology approaches. We present RAC1 as an exemplar and emerging molecule with potential for medical translation.

## Introduction

Mechanotransduction is the event that consists of the conversion of a mechanical signal into a biochemical one (sometimes via further transductions). All our cells are sensitive to mechanical stimuli, implying that all are able to perform this conversion. This competence is crucial in a number of situations, ranging from the very basic mechanisms of early embryonic development ^1^, to the medically relevant spread of metastases^2^. These are two highly divergent variants of the epithelial mesenchymal transition (EMT), type I and III, respectively, which are relevant in experimental regenerative medicine and mechano-pharmacology^3,4^.

Nevertheless, the exploitation of this sensitivity is not part of clinical routine, and, in medical practice only a selected set of specialties make explicit usage of the ability of mechanical stimuli to impinge on yet another EMT (type II) i.e. *wound healing*: dermatology for scarring and burning ^5^, orthopedics for bone repair^6^.

Our interest here is in the connection between mechanical stimuli and inflammation, namely, in the mechanical events whose transduction impacts on the inflammatory function, which plays a crucial role in wound healing. Although numerous pieces of information are available in the literature, their integration is relatively limited^7^. This hampers the development of a systemic overview that is crucial to fully capture the potential and limitations of using mechanical stimuli to modulate inflammation.

One very effective frame to overcome this limitation is the representation of information in a standardized pathway, which exploits the network formalism. This allows to capture, rather than simplify, the complexity, enabling the representation of multiple connections (edges of the network, biological reactions) among entities (nodes of the network, biological molecules).

This concept underpins the design of *in silico* pathways, enabling not only a faithful representation of the complexity of biological interactions, but also the exploitation of computational tools proper to the network theory (systems biology). This approach enables the identification of mathematically relevant nodes, whose biological importance has already been assessed ^8^.

Therefore, building on our earlier work^9^, we have here curated a state-of-the art standard representation of the pathway of mechanotransduction in relation to inflammation. This map is represented in a standardized format to ensure exportability, verifiability and expansion using the Systems Biology Markup Language (SBML^10^).

To the best of our knowledge this offers a unique representation of mechano-transduction *in silico that* evolves f rom our own previous effort and offers an invaluable tool for any further automated and systemic exploration.

After curating this map, we performed two sets of experiments: the first to assess the differential impact of various types mechanical stimuli; the second to highlight how incorporating mechanotransduction within the representation of the inflammatory pathway (innate immune response) let emerge new molecules whose relevance as targets or biomarkers should be explored.

Our experiments suggest that different types of mechanical stimuli elicit a bulk of shared functions, but also show emphasis on distinct molecular functions (as they can be seen in the enrichment and diffusion analysis), and that molecules whose inflammatory relevance is generally disregarded, may become important shall mechanical stimuli be impacting on the system (see Topological analysis and in particular RAC1 and CALM1).

## Material and Methods

### 1 Construction of the Mechanotransduction SBML Pathway

The *mechano core* map^9^ has been updated with new entities from 12 recent (2015-2023) comprehensive review articles^11–22^ and f ive research articles specifically related to the review of the role of the aryl hydrocarbon receptor (AhR^23–27^), which, in addition to its recognized role as a sensor for endo - and xenobiotics, has recently been recognized as having role in the transduction of mechanical signals, particularly those associated with endothelial fluid shear stress and directional migration. Publications have been manually selected using the keywords “mechanotransduction”, “mechanical stimuli”, and “mechanical force”.

The map construction was done by manual curation using CellDesigner.4.3^28^, it contains 8 different stimuli including -beyond the generic (*inflammation*) and experimental inflammatory stimuli (lipopolysaccharide LPS *stress)*-*mechanical force, shear stress, stretch, stiffness, tension*, and *hydrostatic pressure*. All components were updated with the standard MIRIAM (Minimal Information Requested in the Annotation of Models following-up on previously defined and accepted standards ^29^) using NCBI gene IDs for genes, UniProt ID for proteins and ChEBI ID (Chemical Entities of Biological Interest) for small molecules, within the SBML *bqbiol:is* label. Entities that have more than one ID were added to the *bqbiol:hasPart* element according to SBML rules^30^. Multiple proteins or genes with physical interaction are known as *complexes*, while multiple entities without physical interaction are known as *sets*. Sets are further divided in: *DefinedSet* (very general way of grouping molecules based on the fact that they share some common property, they often represent real complexes), CandidateSet (way of grouping molecules when information is incomplete and one of several candidate physical entities is responsible for a particular task , indicating potential alternatives), this latter attributes were added to the *note*, to be fully compliant with Reactome^31^. Finally, the literature pertains to experiments conducted on multiple organisms. This information is stored in the CellDesigner *layer* attribute. This constitutes the complete update of the mechanotransduction pathway in SBML and can be found at Biomodels ID MODEL2406110001 and in the Supplementary Material).

### 2. Construction of three mechano-inflammatory networks

The SBML format enables *systems biology* approaches to the study of mechanotransduction. In particular, pathways represented as complex networks can be investigated in search of relevant nodes or structures, whose role as biomarkers or target is now well assessed. To perform this analysis we used Cytoscape (Version: 3.10.1) with the cy3sbml app ^32^ to import the SBML file.

We first expanded the core mechanotrasduction network described above to include well known sub - pathways. For this, we manually identified 17 nodes whose relevance in inflammation is well known, to extend their neighborhood via Reactome (Table 1). We name this network *Mechano-Union*.

**Table 1:** Subpathways of the Mechanotransduction pathway.

| S.No | Reactome Pathway ID | Reactome Pathway Name |
| --- | --- | --- |
| 1 | R-HSA-171007 | p38MAPK events |
| 2 | R-HSA-195721 | Signaling by WNT |
| 3 | R-HSA-983705 | Calcineurin activates NFAT |
| 4 | R-HSA-2028269 | Signaling by hippo |
| 5 | R-HSA-8878166 | Transcriptional regulation by RUNX2 |
| 6 | R-HSA-9006936 | Signaling by TGFB family members |
| 7 | R-HSA-9013149 | RAC1 GTPase cycle |
| <u>8</u> | R-HSA-5603029 | IkBA variant leads to EDA-ID |
| <u>9</u> | R-HSA-75893 | TNF signaling |
| <u>10</u> | R-HSA-1980143 | Signaling by NOTCH1 |
| <u>11</u> | R-HSA-1489509 | DAG and IP3 signaling |
| <u>12</u> | R-HSA-166016 | Toll Like Receptor 4 (TLR4) Cascade |
| <u>13</u> | R-HSA-112409 | RAF-independent MAPK13 activation |
| <u>14</u> | R-HSA-9818026 | NFE2L2 regulating inflammation associated genes |
| <u>15</u> | R-HSA-8939211 | ESR-mediated signaling |
| <u>16</u> | R-HSA-6783589 | Interleukin-6 family signaling |
| <u>17</u> | R-HSA-194138 | VEGFA-VEGFR2 Pathway |

We also defined a second network, by integrating the *Mechano-Union* to the Reactome Innate immune system (R-HSA-168249), this represents the expansion of the innate immune response to include also mechanical stimuli, in line with the idea of a greater inflammatory pathway able to account for physical transduction^7^. We name this network *Mechano-Innate*.

Finally, our third network was obtained by simply importing the Reactome *Innate immune system* pathway to use it as a baseline for comparison of network analyses (section 2.2). We name this network *Innate*.

All three Cytoscape networks are available as Supplementary material (MechanoUnion.cyjs used in Section Diffusion and MechanoInnate.cyjs, Innate.cyjs used in Section Topological Analysis).

### 3 Diffusion Analysis

Network propagation or diffusion analysis was done using the *Diffusion* App of Cytoscape ^33^ and applied to the *Mechano-Union*. Briefly, this analysis enables us to follow information fluxes (i.e. thermal energy accumulated on some nodes) over time. In a translational perspective, this allows us to explore how signaling flows in the network, i.e. what are the final actuators of initial stimuli. For this reason the analysis was run, setting an amount of energy solely on the node representing a stimulus of interest (one by one), and collecting the amount of energy that reaches the nodes of the network at the end of the simulation, after heat has been transferred to the nodes encountered^34^. Results consist of a ranked list of the network nodes (i.e. with nodes having collected the highest amount of information/energy ranking higher).

This type of output is well suited to perform an *enrichment analysis* which enables the identification of biological functions that are statistically significantly associated to the nodes that preserve energy: in other words, this allows the identification of the biological functions elicited by the stimulus^35^. This was done with R software version 4.0.4 and RStudio version 1.4.1106 by function *gsea*^35^ in the Bioconductor Package *clusterProfiler*, and using the Hallmark gene sets ^36^ as functional references. Results significance is corrected also by the number of tests (i.e. eight stimuli). Given that Hallmark gene sets and Reactome pathways do not use the same standard notation, we performed an automatic names conversion on all SBML pathways before importing them in Cytoscape. The MIRIAM identifier (see Section 1 Construction of the Mechanotransduction SBML Pathway ) enclosed within the SBML’s sub-element “annotation” was extracted and used to query the BioMart R package (version 2.46.3)^37^. In particular we used the “hsapiens_gene_ensembl” and within this set the “uniprot_gn_id”, “entrezgene_id”, “ensembl_gene_id” and “ensembl_transcript_id” columns for the corresponding MIRIAM identifiers (Uniprot, NCBI and Ensembl for genes and transcripts, respectively). These values were used to extract the corresponding species’ name from the “external_gene_name” column, which is then saved in the attribute “name” of each selected species inside the SBML-converted list. In addition to this value, the “name” of each species also contains in brackets the compartment name.

This was applied automatically to simple molecules (i.e. not to sets) to 19 SBML files (Mechanotransduction pathway, 17 pathways in Table 2, and Reactome Innate immune system (R-HSA-168249)).

**Table 2.** Immuno-mechano complexes.

| Node Complex Name | Group |
| --- | --- |
| 2x DDX58 ligand:2x DDX58:2xATP [cytosol] | pathogens |
| Activated TAK complexes [cytosol] |  |
| STING:TBK1:IRF3 [cytoplasmic vesicle membrane] |  |
| TAK1 complex [cytosol] |  |
| UBE2N:UBE2V1 [cytosol] |  |
| WRC:IRSp53/58:RAC1:GTP:PIP3 [plasma membrane] |  |
| CHUK:p-S177,S181-IKBKB:IKBKG [cytosol] |  |
| clustered IgG-Ag:p-FCGRs [plasma membrane] |  |
| clustered IgG-Ag:p-FCGRs:p-6Y-SYK [plasma membrane] |  |
| dsDNA:ZBP1:pS-172-TBK1 [cytosol] |  |
| IKBKG:p-S176,S180-CHUK:p-S177,S181-IKBKB [cytosol] |  |
| Mother filament:branching complex:daughter filament [cytosol] |  |
| NLRP4:DTX4:dsDNA:ZBP1:pS-172-TBK1 [cytosol] |  |
| NLRP4:DTX4:STING:p-S172-TBK1:IRF3 [cytoplasmic vesicle membrane] |  |
| p-S,2T-IRAK4:oligo-MyD88:TIRAP:activated TLR receptor [plasma membrane] |  |
| DAGs [plasma membrane] | lipids |
| PI(3,4,5)P3 [plasma membrane] |  |
| PI(4,5)P2 [plasma membrane] |  |
| PPi [cytosol] |  |
| DAP12 Receptors:p-DAP12:p-6Y-SYK [plasma membrane] | DAP12 |
| p-5Y-LAT:GRB2:SOS1:GADS:p-Y113,Y128,Y145-SLP-76:PLCG:VAV:BTK:PIP3 [plasma membrane] |  |
| p-5Y-LAT:p-SHC1:GRB2:SOS1:GADS:p-Y113,Y128,Y145-SLP-76:PLCG [plasma membrane] |  |
| NFKB1(1-433), NFKB2(1-454):RELA [cytosol] | NFKB |
| NFKB1:MAP3K8:TNIP2 [cytosol] |  |
| p-S32,36-IkB-alpha:NF-kB complex [cytosol] |  |
| Active NIK:p-S176,180-IKKA dimer [cytosol] |  |
| Clustered p:LYN:p-FCERI:IgE:allergen:p-6Y-SYK [plasma membrane] | FCERI |
| p-4S,T404-IRF3,p-S477,S479-IRF7 [cytosol] |  |
| Allergen:p-LYN:p-FCERI:IgE aggregate [plasma membrane] |  |
| p-5Y-LAT:p-SHC1:GRB2:SOS1:GADS:p-Y113,Y128,Y145-SLP-76:PLCG1:VAV [plasma membrane] |  |
| K63polyUb [cytosol] | Innate immune |
| activated TLR4:TRIF:K63polyUb-TRAF3:K63polyUb-TANK:p-TBK1/p-IKBKE [endosome membrane] |  |
| CALM1:4xCa2+ [cytosol] | CALM1 |
| RAC1:GDP [plasma membrane] | RAC1 |

### 4 Network Topological Analysis

Topological network analysis was run in Cytoscape on the *Mechano-Innate* and the *Innate* networks, the rationale being to assess what differences emerge shall we consider “classical” immune response (*Innate* network) or also the mechanotransduction phenomenon (*Mechano-Innate* network) as an entry point to the activation of immunity.

Preprocessing involved the removal of small molecules given their ubiquity, which include: H_2_O, ATP, ADP, GTP, GDP, H^+^, Na^+^ and Ca^2+^, as their high connectivity biases the topological analysis, ranking them very high. Further the corresponding cytoscape.JSON files were exported and the topological analysis was run with Python 3.8 and the *NetworkX* package to create directed graphs, a manipulation necessary to convert the JSON files that have species’ nodes connected to reaction nodes, into a network where species are represented in nodes and reactions in edges. This was achieved as follows: species nodes with identical names were merged and new edges between species nodes connected via the same reaction node were created, to preserve the biological pathway's structure. Finally, on these structures , topological analysis was applied computing four *centrality* measures: *closeness* indicate how close a node is to all other nodes; *betweenness* indicates the highest number of shortest paths (number of edges connecting two nodes); *degree* indicates the number of connected nodes and *eigenvalue* centrality gives a measure of the connections to nodes that have high connections.

Finally, we compared, for the molecules shared between the two networks, the centrality measures. We retained the nodes whose differences were the highest, since this represents molecules whose impact on immunity changes most, shall we consider mechanical stimuli as relevant in triggering an immune response. We finally retained for the final analysis only molecules ranking high across all four measures, as a proxy for robustness. Therefore, this analysis lets emerge the molecules whose centrality changes most when considering mechanotransduction as an integral part of innate immunity, thus identifying molecules whose relevance might be overlooked as inflammatory targets, while they may become relevant when the system is perturbed by mechanical stimuli.

## Results and discussion

### Mechanotransduction Map

Figure 1 shows the complexity of the mechanotransduction map. Major cellular compartments and eight different types of stimuli have been included with their associated transductions.

**Fig. 1.**
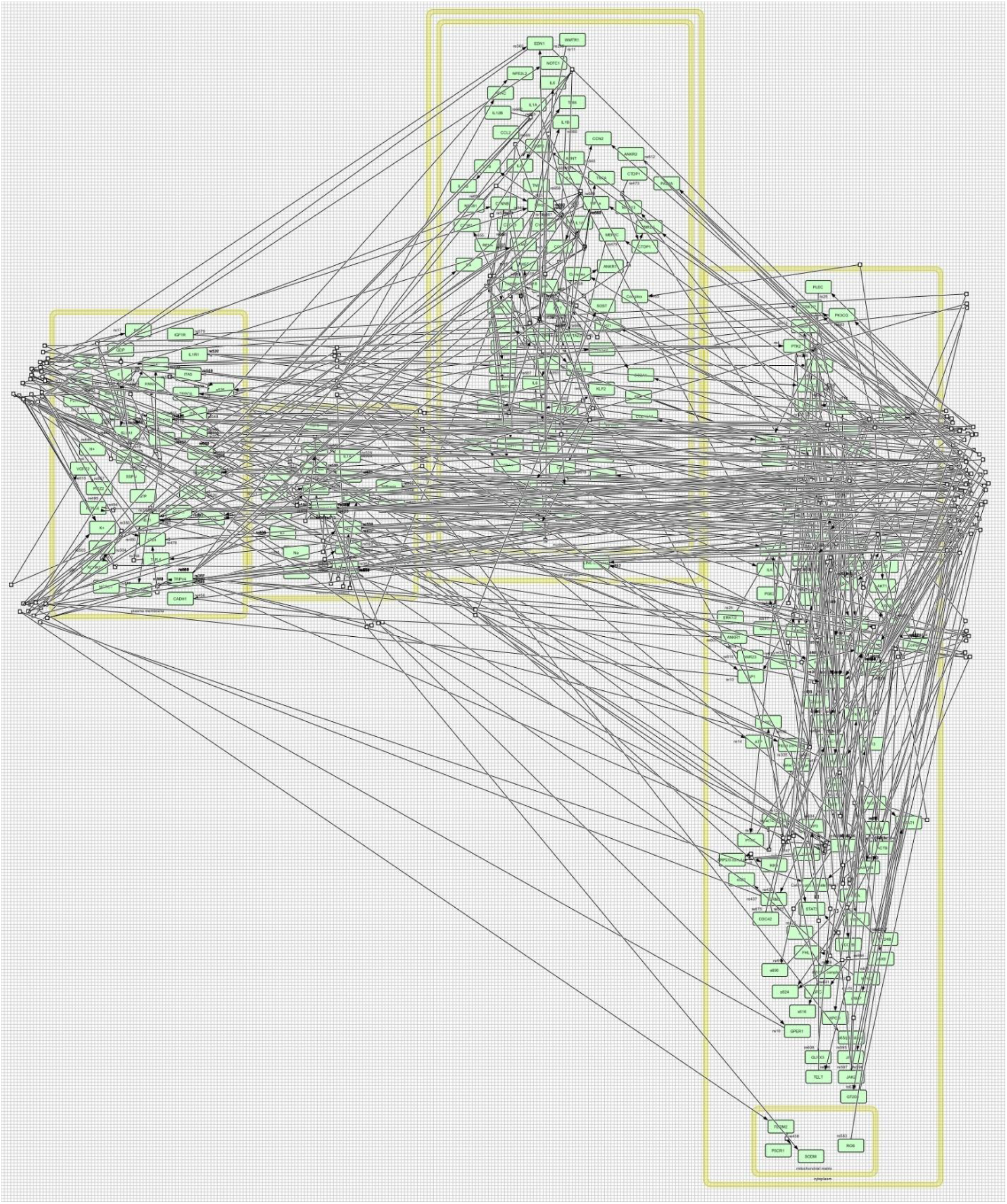
SBML representation of the Mechanotransduction map in SBGN format exported from CellDesigner. This figure is only illustrative of the complexity of the pathway, details can be found in the original SBML in the Supplementary Material or in the Biomodels repository with ID MODEL2406110001

### Diffusion Analysis

The diffusion analysis allows us to gain insight into the steady state (i.e. late) effect of a network perturbation, in our case the different types of mechanical stimuli. In particular, a given amount of energy (heat) is let to flow over time (simulation steps) across the network. According to the specific wiring of the network that will transport a reduced amount of the original energy (consumption) towards connected nodes, while avoiding nodes that are not reached before the exhaustion of this energy.

Results come in the form of a list of all network molecules and their accompanying heat (Supplementary Material, Table S1 Diffusion). This type of information is appropriate for an enrichment analysis, and in particular, *hot* molecules represent high ranking genes (or associated proteins) that can be processed by gene set enrichment analysis^35^ (see Methods) to identify which functions are majorly (and possibly differentially) affected by the stimuli. Fig 2 shows the results of the enrichment analysis (See also Table S2 Supplementary Material), indicating that in general all mechanical stimuli elicit inflammatory and proliferative functions. These functions are broadly in line with the wound healing process (or EMT type II^38^) which proceeds across an early and transient inflammatory phase and eolves towards regeneration (requiring proliferation) and further remodeling.

**Fig. 2.**
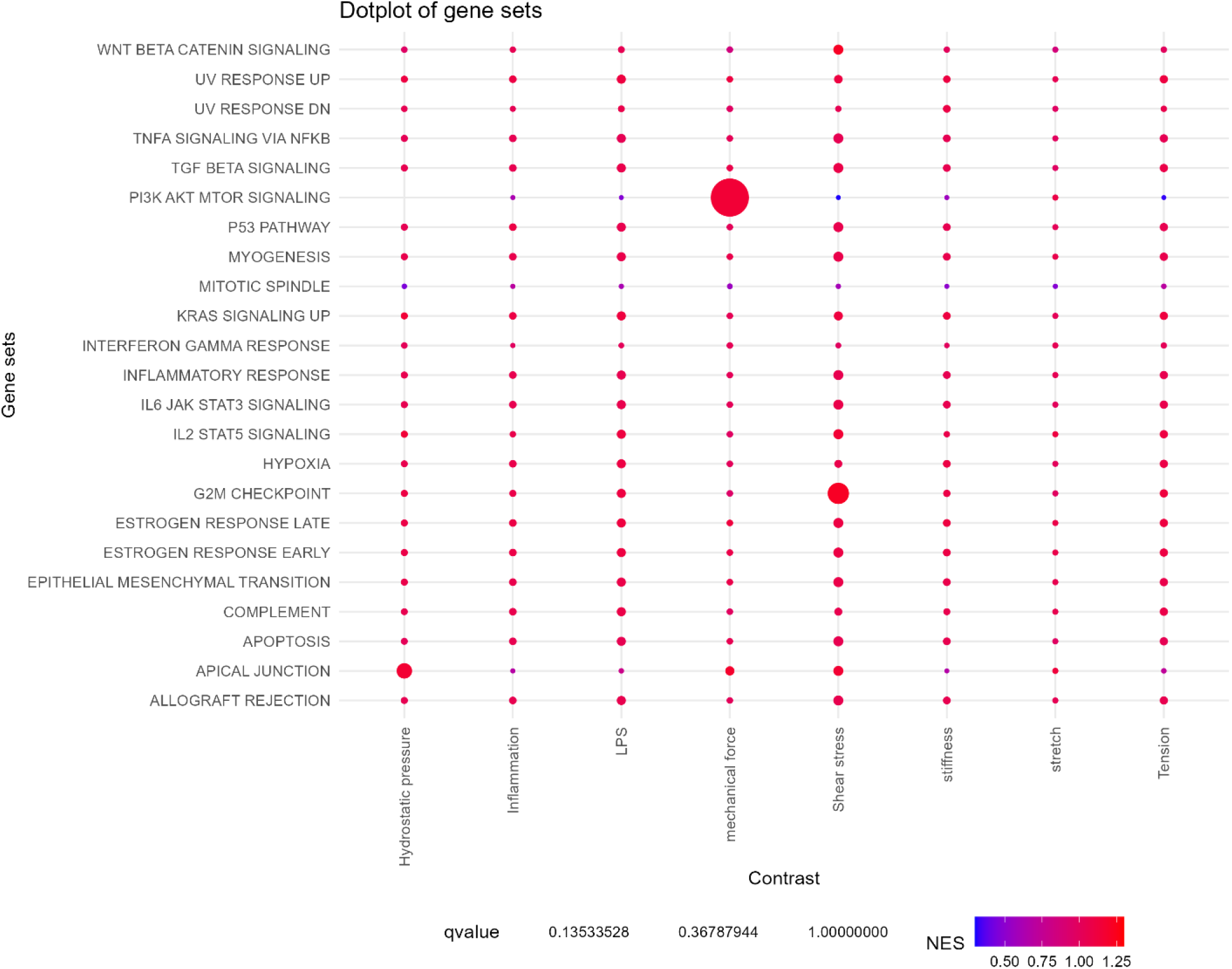
Enrichment analysis (by GSEA) of the network genes ranked by heat, upon diffusion analysis. The functions on the vertical axis correspond to enriched Hallmark functions, while the horizontal axis reports the elicited stimulus. The size of the dots is inversely proportional to the statistical significance of the enrichment.

It is nevertheless interesting to observe that, beyond these common functions, some specific patterns can be observed. Not surprisingly, *inflammation* and *LPS*, at this granularity of modeling, indicate a relatively uniform activation of all the functions identified. Conversely, *mechanical force*, which represents the indistinct mechanical stimulus that was modeled in our previous work^9^ shows a visible enrichment on the tumor associated *mTOR pathway* ^39^, this is in line with the literature that was adopted for that reconstruction, which turned out to have a particular emphasis on EMT type III. This bias was among the motivations to update our network with the current work.

Finally, regarding the specific types of mechanical forces included in our model, we can observe few peculiar patterns. The first is concerned with *hydrostatic pressure* which elicits more specifically *apical junctions* (formation), a results that, if experimentally validated, could expand the information reported from the original literature^27^. The second regards *shear stress*, where the activity concerned with the *GM2 checkpoint* appears to be the most enriched. Interestingly we can also observe that mechanical *stretch* appears to elicit the same overall functions but with lower significance, suggesting, in this sense, a minor efficacy of this type of stimulus.

### Topological Analysis

Topological analysis allows the identification of important nodes, where importance is defined based on the ability of such nodes to connect the network. The most assessed metrics to compute this importance include the *degree*, the *betweenness, eigenvalue* and *stress centrality* and the *eigenvalue*. In our analysis we computed all statistics to obtain a list of molecules whose importance is dramatically different when computing it in the Innate only or in the Mechano-Innate, i.e. including also mechanotransduction. We here discuss the role of these molecules and do so after selecting the more robust, i.e. the ones which change their importance across all four measures (full list and metrics in Table S3, Supplementary Material).

The full list of such molecules and their complexes (34 nodes in total, including mostly complexes) that we name here *immuno-mechano complexes*, is available in Table 2 and Table S4 of the Supplementary Material. We retrieved from Reactome information on the main role of such complexes, and grouped them by six main areas of activity, color coded in Figure 3 (and Table S5 in Supplementary Material): the first pertains directly (and generically) to the response to pathogens. The second has to do with membrane lipids, this is not surprising, as it is well known that mechanical transduction consists of the activation of receptors by physical stress of the membrane and/or the extracellular matrix. Similarly we observe that the third area has to do with well-known key molecules including *Nuclear Factor kappa B*, NFKB^40^, *Transmembrane Immune Signaling Adaptor (TYROBP)* known as DAP12^41^ and interferons (IFN alpha, beta). Finally, we highlight two specific molecules RAC1 and CALM1, whose biological role is known to be crucial in the regulation of ion signaling. While CALM1 mechano-mediated effects are less explored and so its role as drug target^42^, RAC1 (with RhoA) is a well-known key player in mechanotransduction ^43,44^. Its relevance in inflammation is also known, and the modulation of RAC1 in numerous diseases from allergies ^45^ to cancer^46^ is recognized.

From a topological standpoint, the relevance of these molecules appears to be coherently increased across all measures of centrality (in the plasma membrane and in the cytosol, respectively, see Supplementary Material Table S3). This indicates that these molecules, when considering the effect of a mechanical stimulus on the inflammatory state of the system, have a potential to connect to other molecules (eigenvalue, closeness and degree) and more efficiently (betweenness), that is higher than what has been recognized so far, when inflammation was reduced to its more classical innate immune response definition.

What we offer with our investigation is therefore the recommendation to explore the potential of constraining RAC1 (and CALM1) activity with mechanical stimuli, this and similar approaches (see for instance the modulation of Ca^2+^ ion channels by magnetic stimuli^47^), can efficiently complement pharmacological research and drug design.

## Web site address

## List of legends

**Table 1** Reactome pathways completing the Mechanotransduction pathway

**Table 2** List of the 34 Immuno-mechano complexes identified by topological network analysis

