## Supplementary figures and images for "Mechanotransduction and inflammation: an updated comprehensive representation"

### MechanotransductionSBMLPathway_HR.pdf

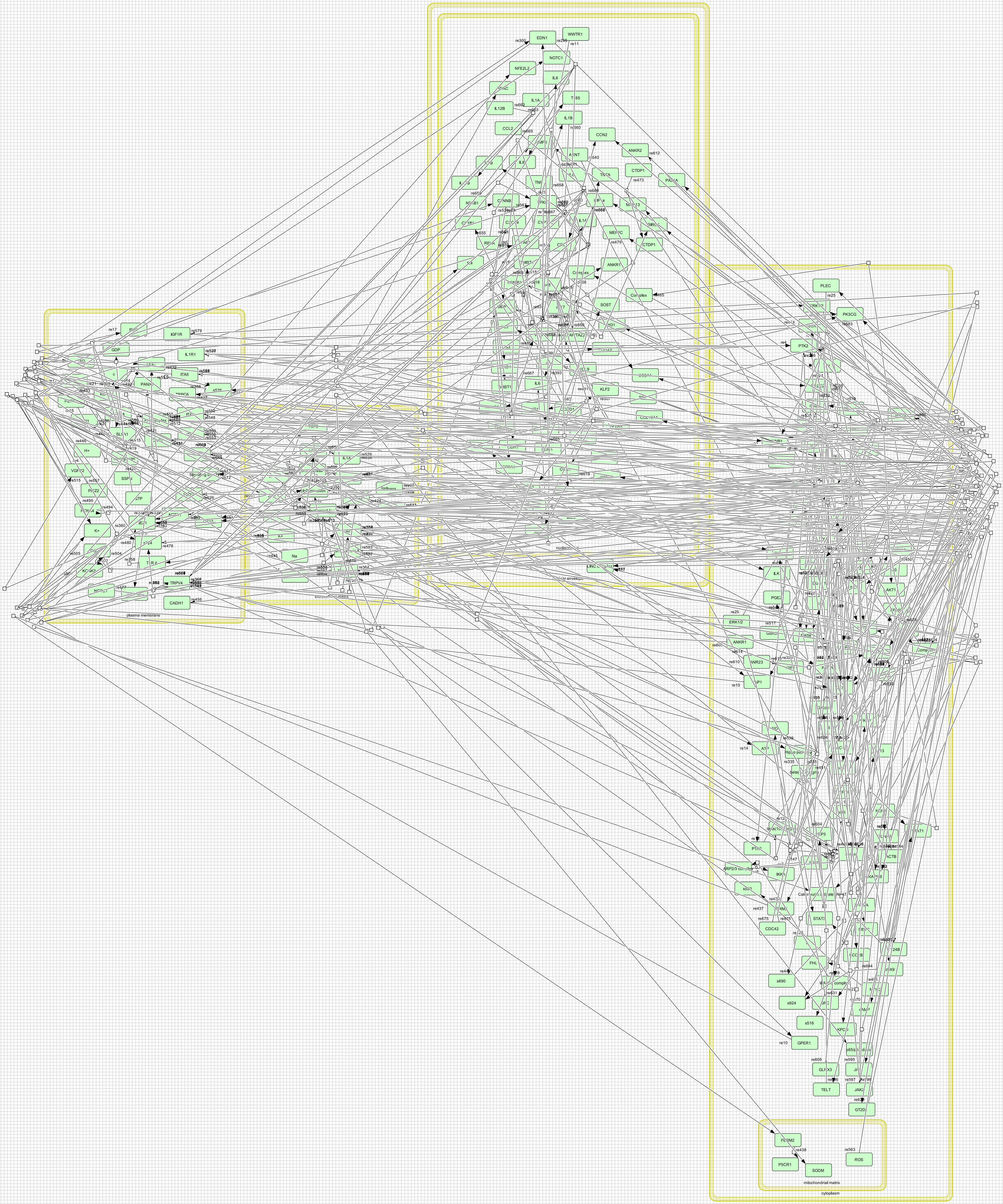
